# A Comparative Study of the Turnover of Multiciliated Cells in the Mouse Trachea, Oviduct and Brain

**DOI:** 10.1101/2019.12.19.882878

**Authors:** Elle C. Roberson, Ngan Kim Tran, Mia J. Konjikusic, Rebecca D. Fitch, Ryan S. Gray, John B. Wallingford

## Abstract

In mammals, multiciliated cells (MCCs) line the lumen of the trachea, oviduct, and brain ventricles, where they drive fluid flow across the epithelium. Each MCC population experiences vastly different local environments that may dictate differences in their lifetime and turnover rates. However, with the exception of MCCs in the trachea, the turnover rates of these multiciliated epithelial populations at extended time scales are not well described. Here, using genetic lineage-labeling techniques we provide a direct comparison of turnover rates of MCCs in these three different tissues. We find that oviduct turnover is similar to that in the airway (∼6 months), while multiciliated ependymal cells turnover more slowly.

## Introduction

The turnover rate of cell populations in different tissues is relevant to our understanding of stem cell biology, cellular response to environment and injury, and cellular function during homeostasis and disease. Multiciliated cells (MCCs) are a terminally differentiated cell type that partially comprise the epithelial lining of the trachea, oviduct, and brain ventricles (Brooks and Wallingford, 2014; Spassky and Meunier, 2017). At their apical surface, MCCs contain dozens of motile cilia, which beat synchronously to drive fluid flow across an epithelium (Brooks and Wallingford, 2014; Spassky and Meunier, 2017). Specifically, MCCs in the trachea are responsible for mucociliary clearance whereby mucus and debris are driven out of the respiratory system, preventing chronic airway infections (Knowles and Boucher, 2002). In the oviducts, MCCs are potentially important for female fertility, although the exact role for these cells is still being defined (Afzelius, 1976; Vanaken et al., 2017). Finally, MCCs in the brain (termed ependymal cells) create local flows of cerebrospinal fluid (CSF) in the ventricular system that are critical for development and homeostasis (Sawamoto et al., 2006).

While MCCs drive fluid flow in all three tissues, each population of MCCs experience vastly different local environments that may affect their function and lifetime. In the trachea, MCCs are exposed to inhaled chemicals like cigarette smoke, environmental pollutants, and airborne pathogens which are known to injure tracheal MCCs (Abdi et al., 1990). Oviduct MCCs are enriched in the infundibulum (the portion of the oviduct closest to the ovary), where they experience repeated ovulations and inflammatory ovulatory fluid (Duffy et al., 2019). Ependymal cells experience contact with the peptides and molecules in the CSF thought to be important for neuroproliferation, differentiation, and homeostasis (Zappaterra and Lehtinen, 2012). Such differences in local environments might be expected to drive variation in MCC turnover time for each cell population.

In addition, each of these multiciliated populations arise from a progenitor either during development or homeostasis. From previous lineage-tracing studies in the trachea, we know that Krt5-positive basal cells are the progenitors of secretory cells, which in turn give rise to MCCs (Rock et al., 2009). Lineage-tracing during embryogenesis demonstrated that radial glial cells are the progenitors of ependymal cells (Spassky et al., 2005). Finally, mouse oviduct MCCs arise from secretory cells, and while the underlying potential progenitor cell in humans is Krt5-positive, the underlying progenitor cell in mice remains unclear (Paik et al., 2012; Ghosh et al., 2017). While each MCC population is known to arise from a progenitor cell, the lifespan of MCCs has only been quantified in trachea (Rawlins and Hogan, 2008), leaving an important gap in our knowledge.

Using genetic lineage tracing, we have performed a comparative analysis of MCC epithelial turnover rates in the trachea, oviduct, and brain of the mouse. As expected, our data concerning trachea MCCs recapitulate the previously published lifespan (Rawlins and Hogan, 2008). We find that the oviduct MCCs – despite a vastly different local environment and an as yet unidentified population of dedicated stem cells – display a lifespan similar to the trachea MCCs. Finally, we demonstrate that ependymal cells are longer lived compared to the trachea and oviduct MCCs.

## Results and Discussion

### Experimental set-up

To determine the turnover rate of MCCs, a genetic pulse-chase experiment was conducted, similar to that previously described for the trachea (Rawlins and Hogan, 2008). We used the *FOXJ1*^*CreER*^ allele which is specifically expressed in MCCs (Rawlins et al., 2007) and the *Rosa26*^*mTmG*^ reporter allele, which upon Cre recombination switches from membrane-TdTomato to membrane-GFP (mGFP) expression (Muzumdar et al., 2007). *FOXJ1*^*CreER*/+^; *Rosa26*^*mTmG*/+^ animals were injected five times over two weeks with a low dose of tamoxifen (tmx) to mosaically label a subset of MCCs with mGFP (Fig. 1A). Littermate control cohorts were injected with corn oil over the same time frame. Animals were euthanized at various time points spanning ∼1 year after the last tmx injection: 1 week to determine the maximum mGFP labeling and then 4 weeks, 12 weeks, 24 weeks, 36 weeks, and 48 weeks to assess turnover (Fig. 1A). All three tissues were dissected from each animal, and the turnover of each population was determined based on loss of lineage labeled, mGFP^+^, MCCs (Fig. 1B-D). As mGFP is enriched in MCC cilia, we used this to quantify the percentage of ciliated luminal surface covered by mGFP^+^ cilia over time (Fig. 1E).

**Figure 1.**
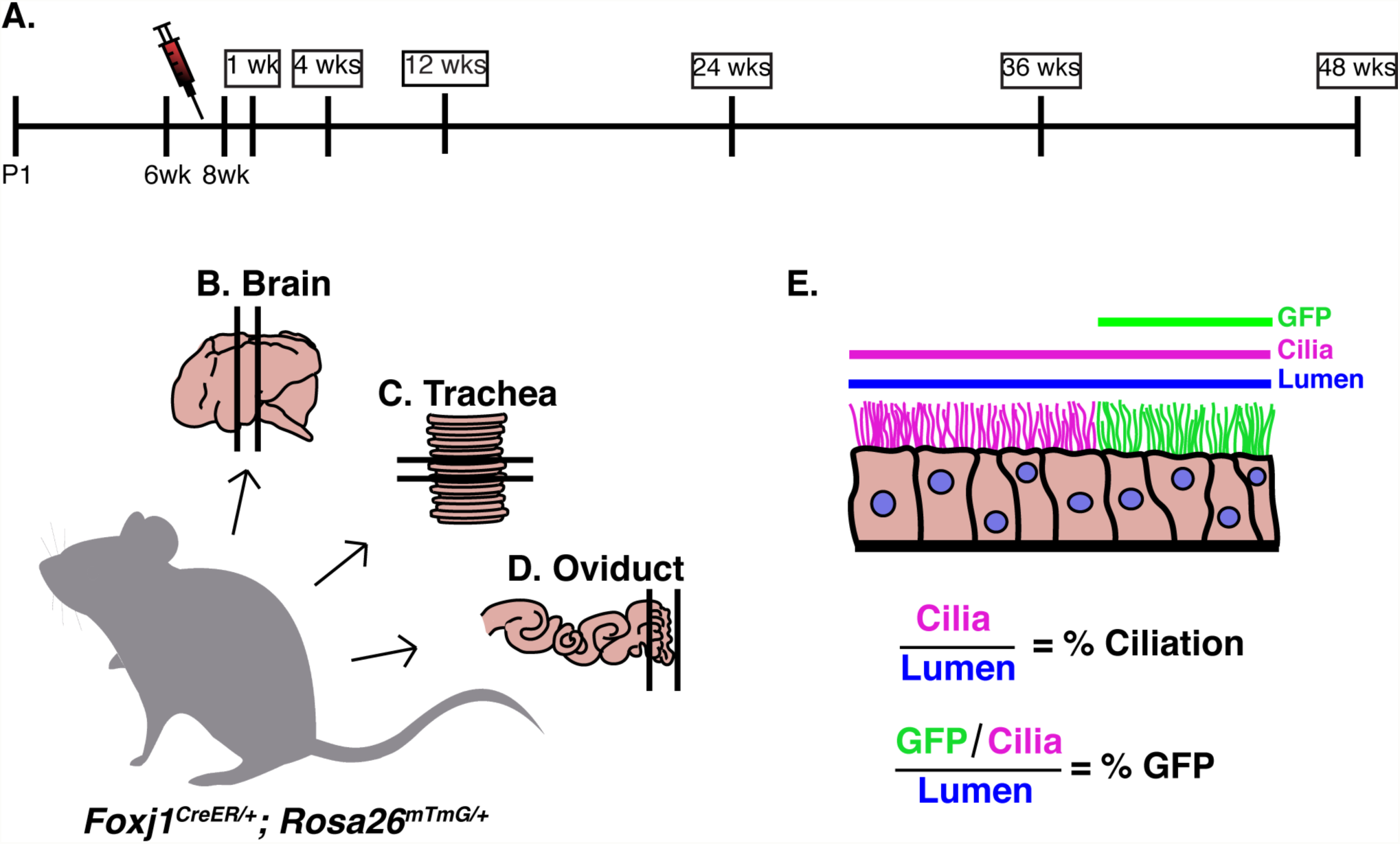
Experimental set-up to perform a comparative analysis of MCC turnover rates. A) Timeline of experiment. *Foxj1*^*CreER*^; *Rosa26*^*mTmG*^ female mice between 6-8 weeks old were injected five times over two weeks with a low dose of tamoxifen (tmx, red needle). Mice were euthanized at various time points after the last tmx injection: 1 week (wk), 4wks, 12wks, 24wks, 36wks, and 48wks. Three tissues were dissected from each animal, the B) brain, C) trachea, and D) oviduct. E) Tissue was analyzed at each timepoint by calculating both the % ciliation and the % GFP-positive cilia lining the lumen. In FIJI, the length of the lumen (blue line), Tub^Ac^ (magenta line), and the GFP-positive cilia (green line) was calculated so that % ciliation and % GFP could be calculated.

### Trachea MCCs

Trachea sections from each animal were probed for acetylated tubulin (Tub^Ac^) to label cilia (Fig. 2A,C) and GFP to mark the genetically labeled MCCs (Fig. 2B,C). At 1 week, ∼38% of cilia were mGFP^+^, while at 48 weeks, ∼2.9% of cilia were mGFP^+^ (Fig. 2A-C, E). Importantly, analysis of Tub^Ac^+ cells revealed that the overall percentage of ciliated cells in the trachea did not change over the course of the experiment (Fig. 2D). From this dataset, we calculated that the turnover rate of tracheal MCCs is 21.18 weeks, or ∼5.3mo (Fig. 2E).

**Figure 2.**
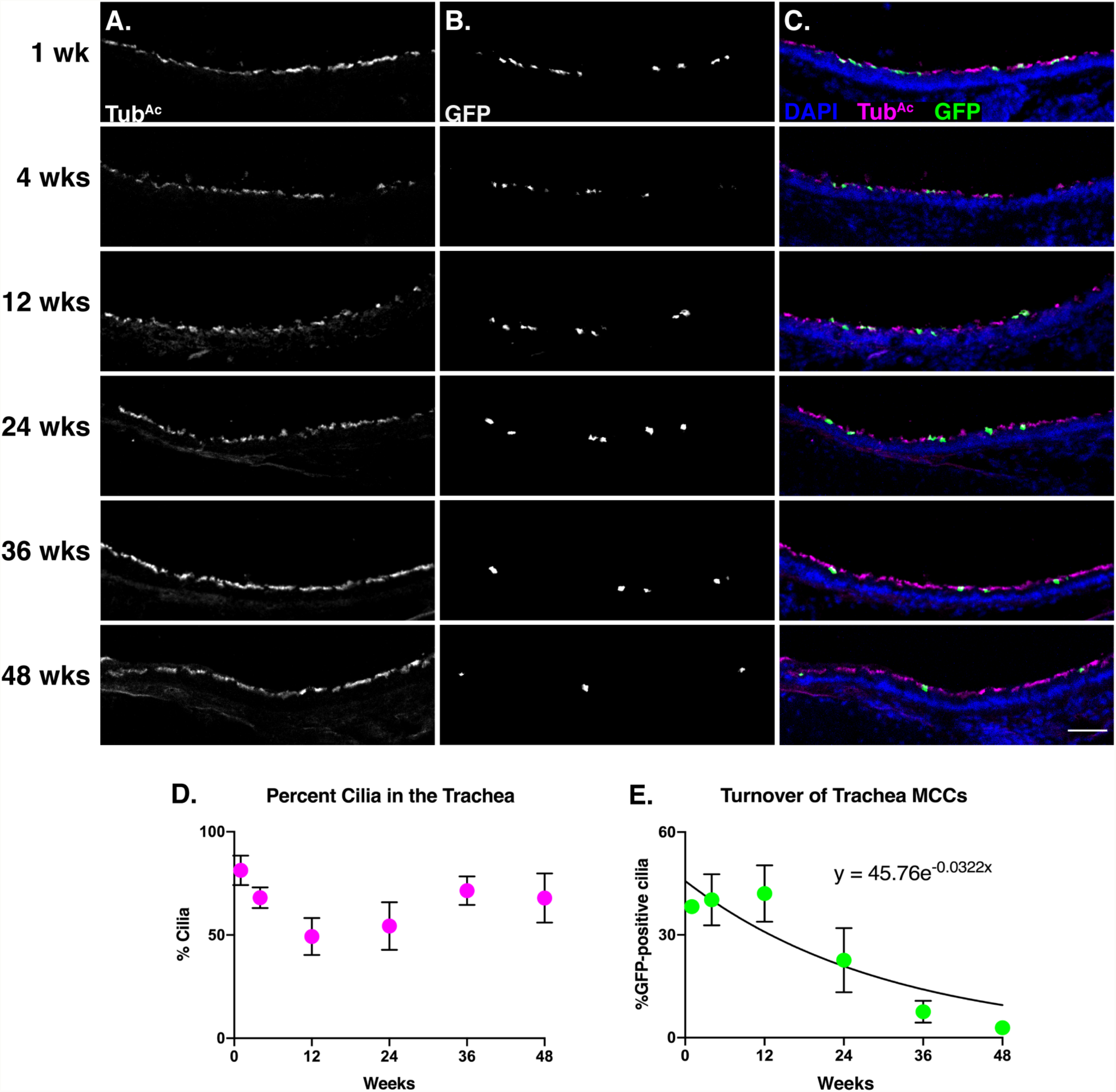
The MCCs of the trachea display a half-life of 5.5 months. Trachea sections from each timepoint were stained for A) cilia (Tub^Ac^, magenta), B) lineage labeled, mGFP^+^, cells (GFP, green), and nuclei (DAPI, blue). Images were taken at 10x. A-B) Grey scale of Tub^Ac^ and GFP to more clearly show both channels. Scale bar = 50μm. C) Merge of cilia, GFP, and DAPI. D) Quantitation of cilia in the trachea at each timepoint shows little change over time. E) Quantitation of mGFP^+^ cells over time, fitted to an exponential decay, whose equation is shown on the graph. This equation was solved to determine that the turnover rate of tracheal MCCs is approximately 5.5 months. All error bars represent the standard deviation.

This result is in accordance with the previously published turnover rate of trachea MCCs as ∼6mo (Rawlins and Hogan, 2008). Importantly, we used a slightly different analysis method, yet reached a very similar conclusion. The previously published dataset had labeled a higher percentage of MCCs (∼72% compared to our ∼38%), because of a higher concentration of injected tmx (Rawlins and Hogan, 2008). As tmx is a weak estrogen analog, our experiment used a lower dose to circumvent the impact of tmx on the female reproductive tract (i.e. the oviduct) and estrous cyclicity (Martin and Middleton, 1978). In addition, the previous work quantified turnover rates by manually counting the number of discrete YFP^+^ MCCs over time based on co-expression of the lineage tracer (YFP) and a cilia marker (β-tubulin). To avoid mis-calling labelled or unlabeled cells, we instead quantified the percentage of ciliated luminal surface covered in mGFP^+^ cilia (Fig. 1E). Nonetheless, we observe that our labeling scheme was effective.

### Oviduct MCCs

Similar to the trachea, oviduct sections were probed with Tub^Ac^ to mark cilia and GFP to mark the lineage-labeled cells (Fig. 3A-C). We focused our quantitation on the anterior of the oviduct, as we found this region to be enriched for lineage-labeled MCCs. At 1 week, ∼32% of cilia are mGFP^+^, while at 48 weeks ∼ 6.8% of cilia are mGFP^+^ (Fig. 3A-C, E). Again, Tub^Ac^ staining revealed that the total percentage of MCCs in the oviduct remained constant across the experiment (Fig. 3D). From this dataset, we determined that the oviduct MCC turnover rate is ∼6mo (Fig. 3E).

**Figure 3.**
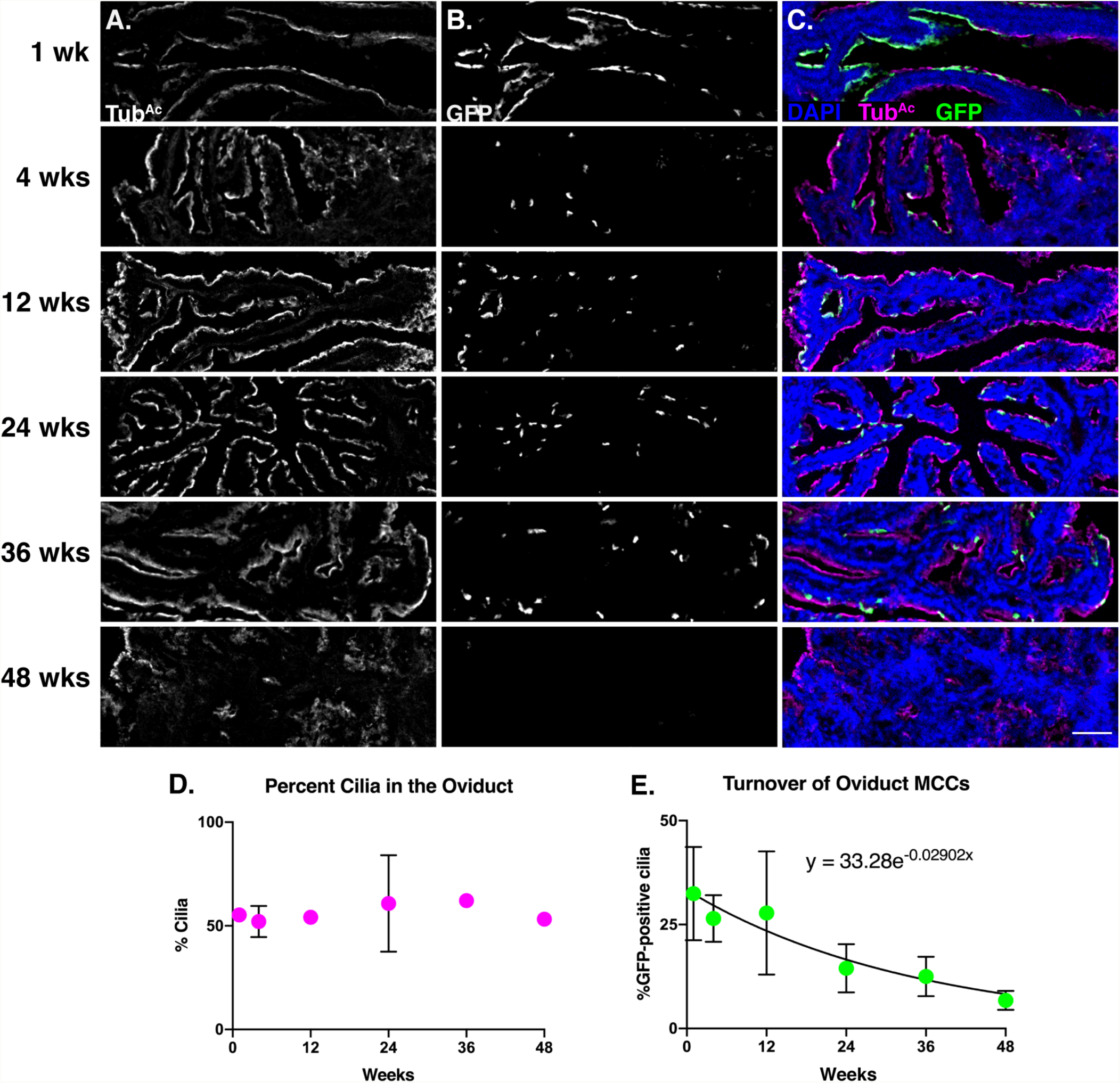
Oviduct MCCs display a half-life of ∼6 months. Oviduct sections from each timepoint were stained for A) cilia (Tub^Ac^, magenta), B) lineage labeled, mGFP^+^, cells (GFP, green), and nuclei (DAPI, blue). Images were taken at 10x. A-B) Grey scale of Tub^Ac^ and GFP to more clearly show both channels. Scale bar = 50μm. C) Merge of cilia, GFP, and DAPI. D) Quantitation of cilia in the oviduct at each timepoint shows little change over time. E) Quantitation of mGFP^+^ cells over time, fitted to an exponential decay, whose equation is shown on the graph This equation was solved to determine that the turnover rate of oviduct MCCs is approximately 6 months. All error bars represent the standard deviation.

Our data suggest that the lifetime of oviduct MCCs is similar to trachea MCCs, which was surprising because the local environment of these two populations is quite different. Trachea MCCs are exposed to various environmental inhalants, while oviduct MCCs are bathed in inflammatory ovulatory fluid every 4-5 days in a mouse (Allen, 1922). The critical implications of ovulatory fluid exposure on the oviduct fimbria has been documented in human studies. In female humans, ovarian epithelial cancer is one of the most prevalent and lethal gynecologic malignancies, and the cell of origin is likely a transformed oviduct fimbrial epithelial cell (Kindelberger et al., 2007; Siegel et al., 2016). Further, a decrease in the number of ovulations – either through pregnancy and lactation, or hormonal contraception – is correlated with a decrease in the risk of ovarian epithelial cancer (Hankinson et al., 1992; Purdie et al., 2003; Tung et al., 2005; Kindelberger et al., 2007; Havrilesky et al., 2013; Troisi et al., 2018). Therefore, recurrent exposure to ovulatory fluid is damaging to oviduct epithelial cells, which increases the risk of ovarian epithelial cancer. In addition, oviduct MCCs are regularly subjected to substantial mechanical perturbation by the passing ova during each estrous cycle. Nonetheless, our data suggest repeated exposure to ovulatory fluid and to ovum transport does not cause oviduct MCCs to turn over more rapidly than trachea MCCs.

This result prompted us to explore the possibility of dedicated stem cells in the mouse oviduct. Mouse oviduct MCCs can arise from secretory cells (Ghosh et al., 2017), though no underlying potential progenitor cell has yet been identified. Interestingly, Krt5+ stem-like cells have been found in human oviduct organoids (Paik et al., 2012), and moreover, Krt5+ basal stem cells give rise to new MCCs in the mouse trachea (Rock et al., 2009). We therefore tested the hypothesis that there may be Krt5+ progenitor cells in the mouse oviduct. Using both a *Krt5*::*GFP* allele (Schoch et al., 2004) as well as lineage labeling with *Krt5*^*CreER*^; *Rosa26*^*mTmG*/+^ (Van Keymeulen et al., 2011), we found no evidence for *Krt5* labeling any cell type of the oviduct (Fig. 4A and C). This result was not due to a failure of the genetic tools, as we did observe labeled *Krt5*-positive basal cells in the trachea as expected (Fig. 4B and D). This study demonstrates that Krt5-positive cells are unlikely to be progenitor cells in the mouse oviduct, and highlights the need for future studies to investigate cell-type development and differentiation in the mouse oviduct. The majority of the developmental lineage of MCCs is derived from work in the trachea (Schoch et al., 2004; Rawlins et al., 2007; Rock et al., 2009), yet this work suggests that the oviduct MCCs differentiate along a different lineage than trachea MCCs, despite their similar turnover times.

**Figure 4.**
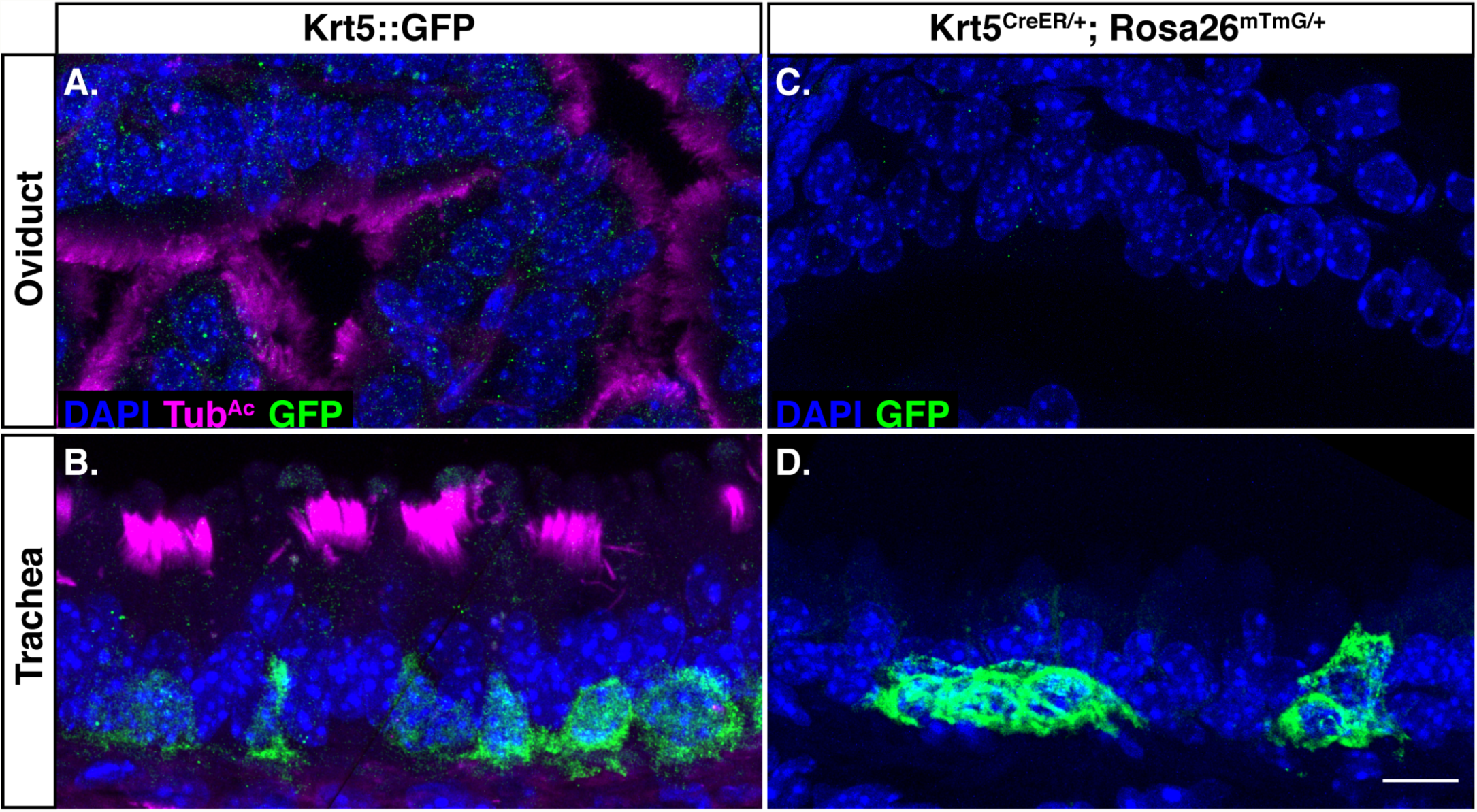
Keratin 5 is not a progenitor marker in the oviduct. *Krt5*::*GFP* females were dissected, and sections of A) oviduct, and B) trachea were probed for nuclei (DAPI, blue), cilia (Tub^Ac^, magenta), and GFP (green). A) The oviduct does not have Krt5-positive cells. B) The trachea acts as a positive control, and does have Krt5-positive basal cells. Sections of C) oviduct, and D) trachea from *Krt5*^*CreER*^; *Rosa26*^*mTmG*/+^ females were probed for nuclei (DAPI, blue), and lineage-labeled cells (GFP, green). C) The oviduct does not have Krt5-positive cells. D) The trachea does display Krt5-positive lineage-labeled basal cells, as expected. Scale bar = 10μm.

### Ependymal cells

Brain sections were probed with Tub^Ac^ to label cilia and GFP to mark the lineage-labeled cells (Fig. 5A-C). In general, our labeling scheme led to much higher proportion of GFP+ ependymal cells compared to MCCs in the other tissues (1wk timepoint in Fig. 5E compared to 1wk timepoint in Fig. 2E and 3E). At 1 week, ∼78% of cilia are mGFP^+^, while at 48 weeks, ∼56% of cilia are mGFP^+^ (Fig. 5A-C, and E). The turnover rate of the ependymal cells was slower than the two other tissues, with only a 25% decrease in GFP+ labeled cell populations over the course of the entire experiment (Fig. 5E). Again, Tub^Ac^ staining revealed that the overall percentage of MCCs in the brain remained constant across the experiment (Fig. 5D). From this dataset, we calculated the half-life of the ependymal cell population to be ∼83 weeks, or roughly a year and a half, which equates to the average lifetime of a normal mouse lab strain (Fig. 5E) (Comfort, 1959).

**Figure 5.**
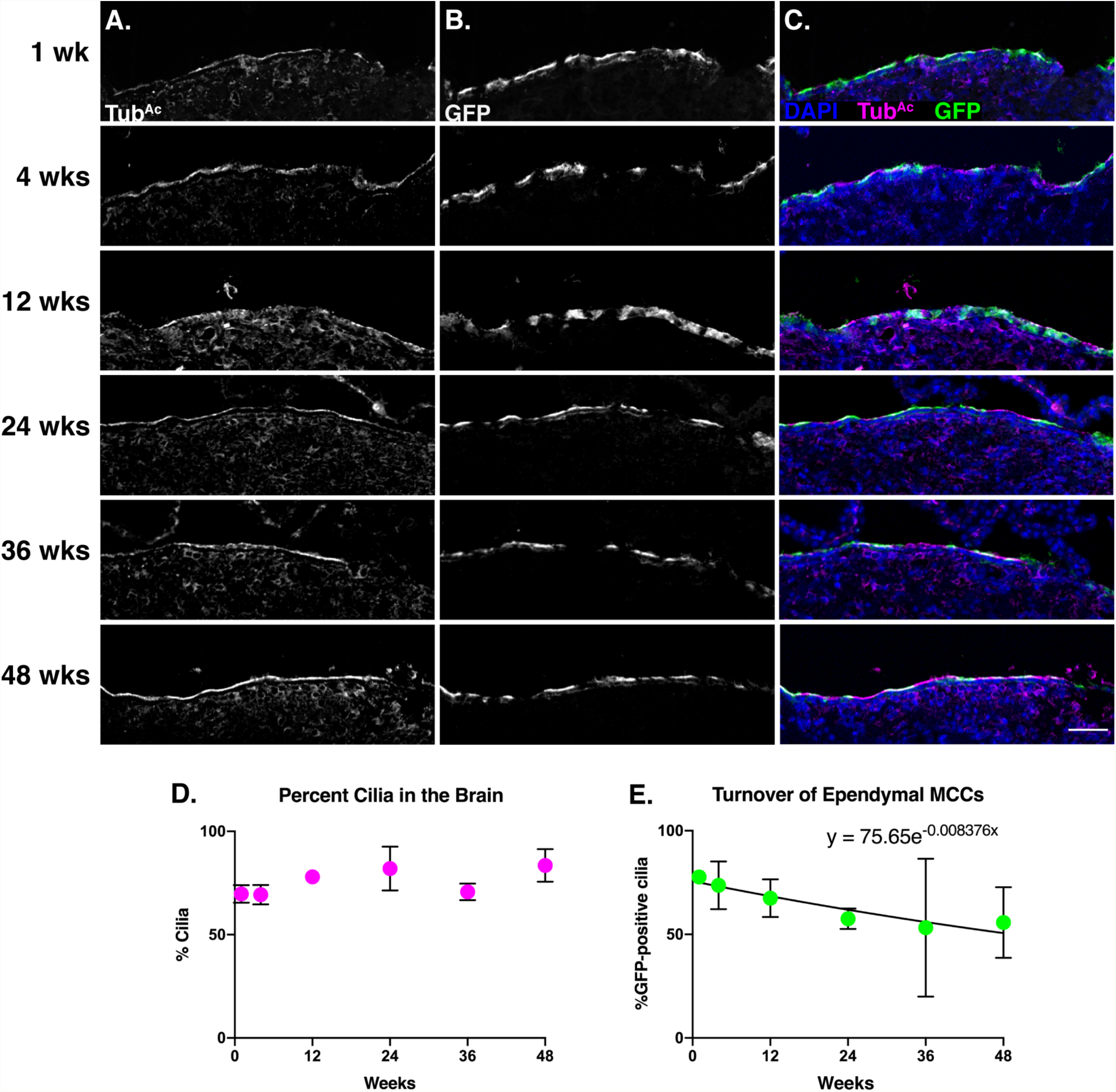
Ependymal cells display a half-life of ∼21 months. Brain sections from each timepoint were stained for A) cilia (Tub^Ac^, magenta), B) lineage labeled, mGFP^+^, cells (GFP, green), and nuclei (DAPI, blue). Images were taken at 10x. A-B) Grey scale of Tub^Ac^ and GFP to more clearly show both channels. Scale bar = 50μm. C) Merge of cilia, GFP, and DAPI. D) Quantitation of cilia in the brain at each timepoint shows little change over time. E) Quantitation of mGFP^+^ cells over time, fitted to an exponential decay, whose equation is shown on the graph. This equation was solved to determine that the turnover rate of ependymal cells is approximately 21 months. All error bars represent the standard deviation.

In contrast to the findings here, a previous study indicated that ependymal cells are post-mitotic and likely born only once during development (Spassky et al., 2005). It is important to note, however, that the previous study focused on a much shorter time frame than our experiment: they labelled mitotically active cells *in utero* and then assessed ependymal cell development by 2mo of age. By contrast, our dataset labels MCCs at 6-8 weeks of age and we collected data for an additional 48 weeks. Because our data spans a much longer time scale, we were able to observe a slight decrease in GFP+ cilia while the percent ciliation of the ventricle stayed approximately the same (Fig. 5D-E). These data suggest, then, that ependymal cells do turnover in the mouse brain - albeit at a much slower rate compared to the trachea and oviduct MCCs. Thus, our data suggest that ependymal cells can be homeostatically replaced, warranting deeper investigation with regards to injury or infection of the ventricular system.

## Conclusion

In conclusion, we have shown that the oviduct and trachea MCCs turnover at approximately the rate (about 6mo), whereas the ependymal cell population turns over significantly slower (about 18mo). These data expand our understanding of the homeostasis of MCC populations, and how different local environments contribute to that homeostasis. The work sets the stage for further understanding how different MCC population lifespans change in response to disease, injury, infection, pregnancy, and infertility.

## Experimental Procedures

### Mice

*FOXJ1*^*CreER2T*^ (Rawlins et al., 2007), *Krt5*^*CreER*^ (Van Keymeulen et al., 2011), *Krt5*::*GFP* (Schoch et al., 2004), and *Rosa26*^*mTmG*^ (Muzumdar et al., 2007) animals were maintained on a mixed background (Swiss Webster and C57BL/6). Mice were housed in individually ventilated cages in a pathogen-free facility with continuous food and water, with a controlled light cycle (light from 7am-7pm). 6-8-week-old female *FOXJ1CreER*^*2T*^; *Rosa26*^*mTmG*/+^ were used to perform a comparative approach of the turnover rates of the trachea, oviduct, and brain multiciliated cells. A cohort of 36 mice were given 5 intraperitoneal (IP) injections of tamoxifen (tmx, Sigma) dissolved in corn oil at the dose of 1mg tmx/40g mouse over 2 weeks (Rawlins and Hogan, 2008). A control cohort of 21 mice were given 5 IP injections of corn oil over the same time frame. Mice in groups of at least 3 were humanely euthanized by extended CO_2_ exposure at 1 week, 4 weeks, 12 weeks, 24 weeks, 36 weeks, and 48 weeks after the final tmx injection (Fig. 1).

6-8 week-old female *Krt5*::*GFP* and *Krt5*^*CreER*^; *Rosa26*^*mTmG*/+^ were used to determine if there are Krt5-positive cells in the oviduct, using trachea as a positive control. Tissue from *Krt5*::*GFP* animals was obtained from Dr. Jeremy Reiter’s lab at UCSF. *Krt5*^*CreER*^; *Rosa26*^*mTmG*^ females were injected once with tmx (1mg tmx/40g mouse) and then euthanized 1wk post-tmx injection. All animal experiments were approved by the University of Texas at Austin Institutional Animal Care and Use Committee.

### Tissue processing & immunofluorescence

Trachea, oviducts, and brains were dissected from each animal and fixed in 4% paraformaldehyde (Electron Microscopy Sciences) overnight at 4°C. Fixed tissues were washed in PBS, and incubated in 30% sucrose overnight at 37°C. Tissues were washed briefly in NEG-50 Frozen Section Medium (ThermoFisher) and then were embedded in NEG-50 in an ethanol/dry ice bath (oviducts & trachea) or allowed to freeze at - 80°C (brains) and stored at −20°C or −80°C. 12μm frozen sections were cut on a cryostat (Leica) and dried overnight at RT. Frozen sections were stored at −20°C.

Tissue sections were washed in PBS + 0.1% Tween20 (PBST) three times to remove NEG-50. Antigen retrieval was performed on brain sections using 0.1 M citrate buffer (pH6.8): citrate buffer was heated to boiling, slides were added to the buffer and allowed to cool for 1hr. Tissues were then blocked for at least 30min at RT with 5-10% normal goat serum + PBS (block buffer). Primary antibodies for GFP (chicken anti-GFP, 1:500 dilution, Abcam, ab13970) and cilia (mouse anti-acetylated tubulin, 1:1000 dilution, Sigma, cat# 6-11B-1) were diluted in block buffer and incubated on slides for 2hr at RT or overnight at 4°C. After washing three times with PBST, tissue was incubated with Alexa-Fluor coupled secondary antibodies (goat anti-chicken 488, goat anti-mouse 647, goat anti-rat 647, 1:1000 dilution, ThermoFisher) and DAPI (1:1000 dilution, ThermoFisher) for at least 30min at RT. After washing at least 3 times with PBST, slides were mounted with Prolong Gold (ThermoFisher).

The oviduct and trachea were imaged for quantitation and publication on a Zeiss LSM700 point scanning confocal microscope. For quantitation, brain tissue was imaged with a CSU-W1 spinning disk Nikon confocal, and publication images were captured with the Zeiss LSM700 point scanning confocal.

### Quantitation

Two to four sections spaced at least 100μm apart were analyzed for GFP^+^ ciliated cells. For the oviduct, we selected sections that contained the most anterior portion due to the enrichment of ciliated cells. For the brain, we selected sections that contained the dorsal third ventricle and lateral ventricles surrounding the anterior hippocampus. As membrane-GFP (from the *Rosa26*^*mTmG*^ transgene) is strongly enriched in the ciliary membrane, we used it to quantify the percentage of luminal surface covered by GFP^+^ cilia. In FIJI (FIJI is Just ImageJ) the lumen of each tissue was traced based on DAPI staining and the length was measured. Similarly, the ciliated surface of the lumen was traced and measured based on acetylated tubulin. Finally, the GFP^+^ cilia were traced and measured along the lumen. From these length measurements, the %GFP^+^ ciliated surface is calculated for each animal at each timepoint. Using GraphPad Prism 8, each dataset was fitted with one-phase exponential decay curves (where the plateau was constrained to 0), which were solved to determine the half-life of MCCs in each tissue.

## Acknowledgements

ECR is supported by the NICHD (F32HD095618). NKT was supported by the TIDES Summer Fellowship Program from the University of Texas at Austin. MK is supported by Provost Graduate Excellence Fellowship through the Institute of Cell and Molecular Biology at the University of Texas at Austin. RSG is supported by the NIAMS (AR072009). JBW is supported by the NICHD (HD085901) and the NHLBI (HL117164). We thank Drs. Scott Randell, Jeremy Reiter and Semil Choksi for generously sharing trachea and oviduct tissue from *Krt5::GFP* female mice.

